# Instructor Perspectives on Challenging Topics in Evolution Education and Their Implications for Game-Based Learning

**DOI:** 10.64898/2026.07.06.736808

**Authors:** Josie Otto, Lauren Goulet, Brooke Kopack Ware, Hannah Lowry, Aeryn-Elayna Miller, Jake D. Botello, Jenna E. Pruett, Abby Beatty

## Abstract

Evolution is a foundational framework for understanding biology, yet it remains challenging to teach and learn. Game-based learning may offer one way to support evolution instruction by helping students visualize abstract, dynamic, and difficult-to-observe processes. In this study, we interviewed undergraduate biology instructors to examine how they evaluated video games as potential tools for evolution education, including which topics they perceived as most challenging for students. We found that instructors were broadly open to using video games for evolution instruction, particularly when games could support population-level reasoning, evolutionary mechanisms, speciation and phylogeny, quantitative reasoning, and long time scales. Instructors also emphasized that games must be scientifically accurate, accessible, and aligned with course learning goals. This study contributes an instructor-centered perspective to evolution education and game-based learning research by identifying how instructors connect persistent student learning challenges with potential design priorities for educational video games.

**Graphical Abstract:** 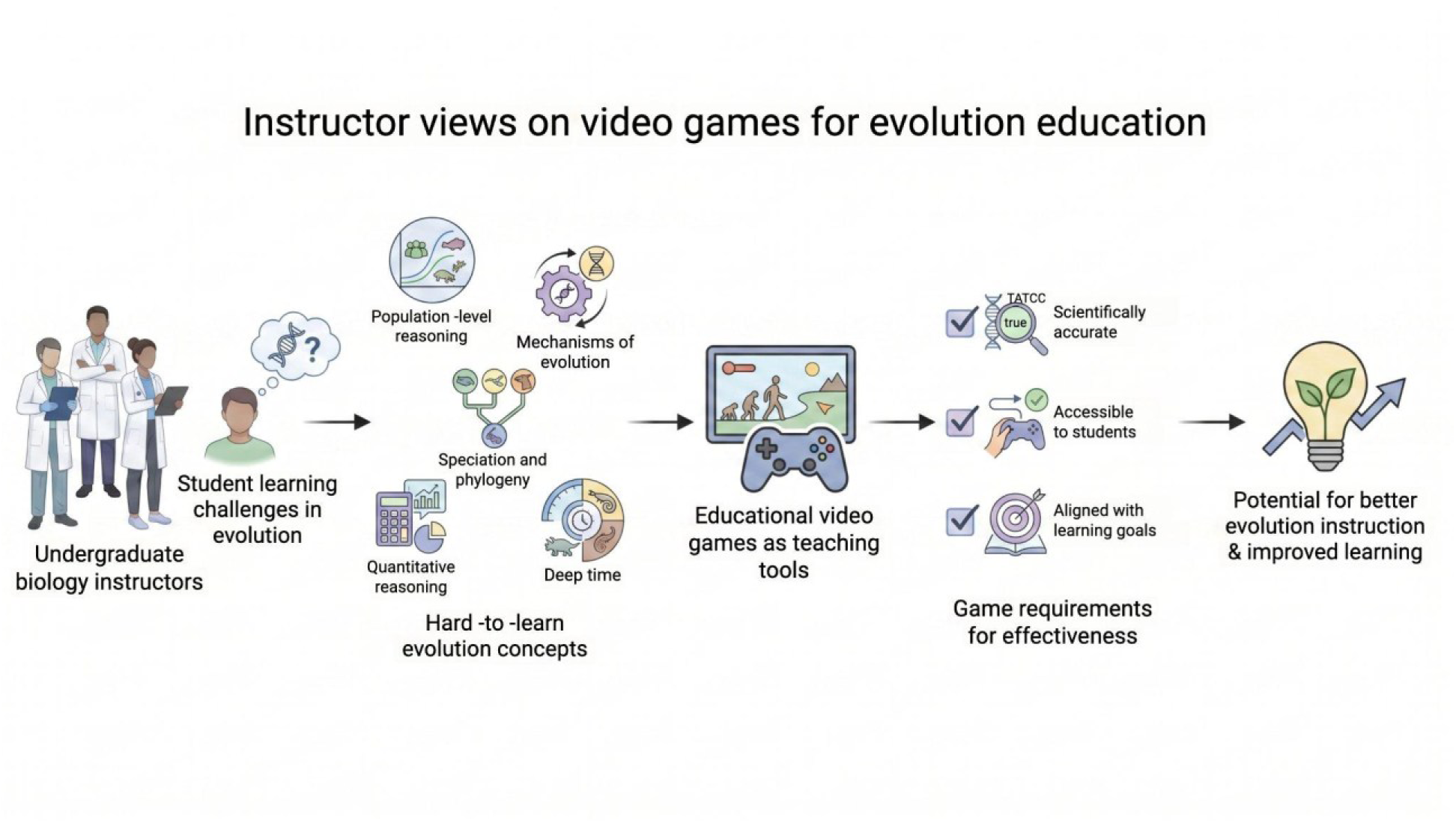

## INTRODUCTION

Evolution is a foundational framework for understanding biology (Dobzhansky, 1964). To understand how biological systems vary, change, and diversify over time, students must also understand the evolutionary processes that shape life across scales. For this reason, evolution has been established as a core concept in undergraduate biology instruction and a focus of national biology education reform (American Association for the Advancement of Science, 2011; Brownell et al., 2014). Yet, evolution remains difficult for many students to understand and accept (Barnes et al., 2024, 2022; Furrow and Hsu, 2019; Misheva et al., 2026). One reason is that many of the processes most central to evolution are not directly observable in everyday classroom settings. As a result, evolution is both essential and unusually challenging to teach and learn.

The difficulty of observing evolutionary processes directly suggests a need for experiential learning tools that allow students to engage with these concepts more concretely. Science educators have long used models, simulations, and other visual tools to help students reason about difficult-to-see processes (Gilbert and Osborne, 1980; Rutten et al., 2012). Interactive simulations are now common in STEM education because they enable students to explore complex phenomena that are difficult to observe directly, including processes that are invisible, occur over long time scales, involve numerous interacting components, or are impractical to recreate in traditional classroom environments (Al-Elq, 2010). For example, PhET simulations have been widely used to support conceptual learning by giving students interactive, visual representations of scientific phenomena (Harahap et al., 2025), including natural selection (Donald, 2025; Su et al., 2021). Simulation-based instruction is also used in biomedical education for clinical skills training by recreating authentic clinical environments and complex procedures (Dankbaar et al., 2016). The use of simulations expanded rapidly during the COVID-19 pandemic, when remote instruction and limited access to in-person laboratories increased demand for digital tools that could support students’ engagement with scientific processes outside traditional lab settings (Owolabi et al., 2025). Their use has continued to grow post-pandemic as access to technology increases and as digital learning tools become easier to develop, distribute, and integrate into instruction. This expanding landscape includes not just simulations, but also other interactive tools that use game-like or game-based design to support student learning.

Simulations belong to a broader family of digital learning tools, but they are not interchangeable with related approaches such as video games, serious games, or gamification (Table 1). In this context, video games refer to full digital games in which players interact with a rule-based environment to pursue goals, make decisions, and experience consequences through play. When full games are designed or used for instructional purposes, they are often discussed as game-based learning (GBL), serious games, or both. Although these terms are not always used consistently, GBL generally emphasizes the use of gameplay to support defined learning outcomes, whereas serious games often refers to games designed for purposes beyond entertainment, including education, training, or skill development (Plass et al., 2015; Ratinho and Martins, 2023; Shaffer et al., 2005). In both cases, the game itself functions as the learning environment rather than as a reward structure layered onto existing instruction. Through gameplay, students may act as decision-makers within a system, exercise agency, test strategies, receive real-time feedback, and revise their actions based on consequences (Nkhoma et al., 2014; Snow et al., 2015). By contrast, gamification refers to “the use of game elements in non-game contexts”, such as adding points, badges, rankings, or levels to a class activity that is not itself a game (Gini et al., 2025). In this study, we focus specifically on video games as a form of GBL, rather than on gamification alone.

**Table 1.** Key terms used to describe games and game-like learning tools.

| **Term** | **Core Idea** | **Is the activity itself a game?** |
| --- | --- | --- |
| Video game | A full digital game with rules, goals, decisions, and consequences | Yes |
| Game-based learning | Using a game to support learning outcomes | Usually yes |
| Serious game | A game designed for education, training, or another non-entertainment purpose | Yes |
| Simulation | An interactive model of a system or process | Not always |
| Gamification | Adding game elements to a non-game activity | No |

Relatively little work has focused on evolution-focused video games for undergraduate biology education. Existing research on evolution games has more often focused on pre-college students, informal learning, or the broad representation of evolution in digital games rather than the design of games for undergraduate evolution instruction (Adams et al., 2011; Bean et al., 2010; Heeter and Starr, 2012; Leith et al., 2016). For example, Leith and colleagues (2016) analyzed how evolution through natural selection was represented in two types of digital games: publicly available games that portrayed evolution and games designed specifically to teach or study evolution. In both sets of games, they examined whether gameplay represented three core principles of natural selection: phenotypic variation, differential fitness, and heritability. They found that games varied widely in whether they represented one, two, or three principles, and that games designed for instruction were often difficult to identify through standard searches. Other work suggests that games do not need to be scientifically perfect to be instructionally useful. In a case study of an upper-level evolution course, SPORE was used as a supplemental teaching tool even though the game represented evolution inaccurately in several ways (Poli et al., 2012). Students who participated in gameplay reported spending more time with course material and scored higher on subsequent exams and in the course overall. However, the game was used alongside instructor-led discussions and writing assignments that asked students to identify and critique scientific inaccuracies. These studies suggest that evolution video games have instructional potential, but scientific accuracy, accessibility, and instructor guidance remain important concerns. As a result, there is limited guidance on how evolution-focused video games should be designed for undergraduate biology classrooms or how instructors evaluate their usefulness.

Instructor perspectives are therefore important for the design of evolution-based educational games. Prior reviews suggest that games and simulations can support student learning, but their effectiveness depends on how they are designed, integrated into a course, and supported by instructors (Clark et al., 2016; Vlachopoulos and Makri, 2017). Instructors have direct knowledge of the concepts students find difficult, the concerns that students bring to evolution instruction, and the tools that are likely to work in their own classrooms. Although prior research has documented barriers to learning and accepting evolution, less is known about how instructors evaluate video games as potential tools for evolution education. Additionally, game developers seeking to produce new games to improve student outcomes may require highly specific guidance to identify areas in which students most struggle. In this study, we address this gap by examining how undergraduate biology instructors across institution types and geographic regions describe the feasibility and utility of evolution-focused game-based learning. We sought to answer the following questions:

1. What conceptual and affective challenges do instructors describe when teaching undergraduate evolution?
2. Which evolutionary concepts do instructors view as well suited to game-based learning?
3. How do instructors’ perceptions of student learning challenges shape their views of video games as instructional tools?

## METHODS

### Data Collection

Participants were recruited through direct email invitations and a national STEM education listserv. Email invitations (n=27) were sent to biology instructors identified through departmental websites and professional networks, and an additional recruitment announcement was distributed via the SABER (Society for the Advancement of Biology Education Research) listserv. The screening survey (Supplemental A) remained open for ten days, during which 39 instructors completed the survey. The survey collected demographic, institutional, and instructional information relevant to participant selection, including institution type and region, course characteristics, years of teaching experience, and instructional appointment or rank (e.g., adjunct, tenure-track, tenured). Demographic characteristics of all survey respondents are reported in Supplemental B.

After the survey closed, respondents were screened using a purposive sampling approach aimed at maximizing diversity across instructor demographics, course types, and institutions. From the pool of 40 respondents, 10 instructors were selected for participation in semi-structured interviews (Table 2). Interviews were conducted remotely using Google Meet over a three-week period and lasted approximately 60-90 minutes. Interview questions focused on instructors’ experiences teaching evolution, instructional goals, and perspectives on game-based learning (see Supplemental C for complete Interview Protocol). All interviews were audio-recorded with Otter.ai and auto-transcribed, then checked and corrected against the audio to ensure verbatim transcripts for analysis. Interview participants received $100 compensation for their time and were invited to participate in the subsequent implementation phase of the video game.

**Table 2.**
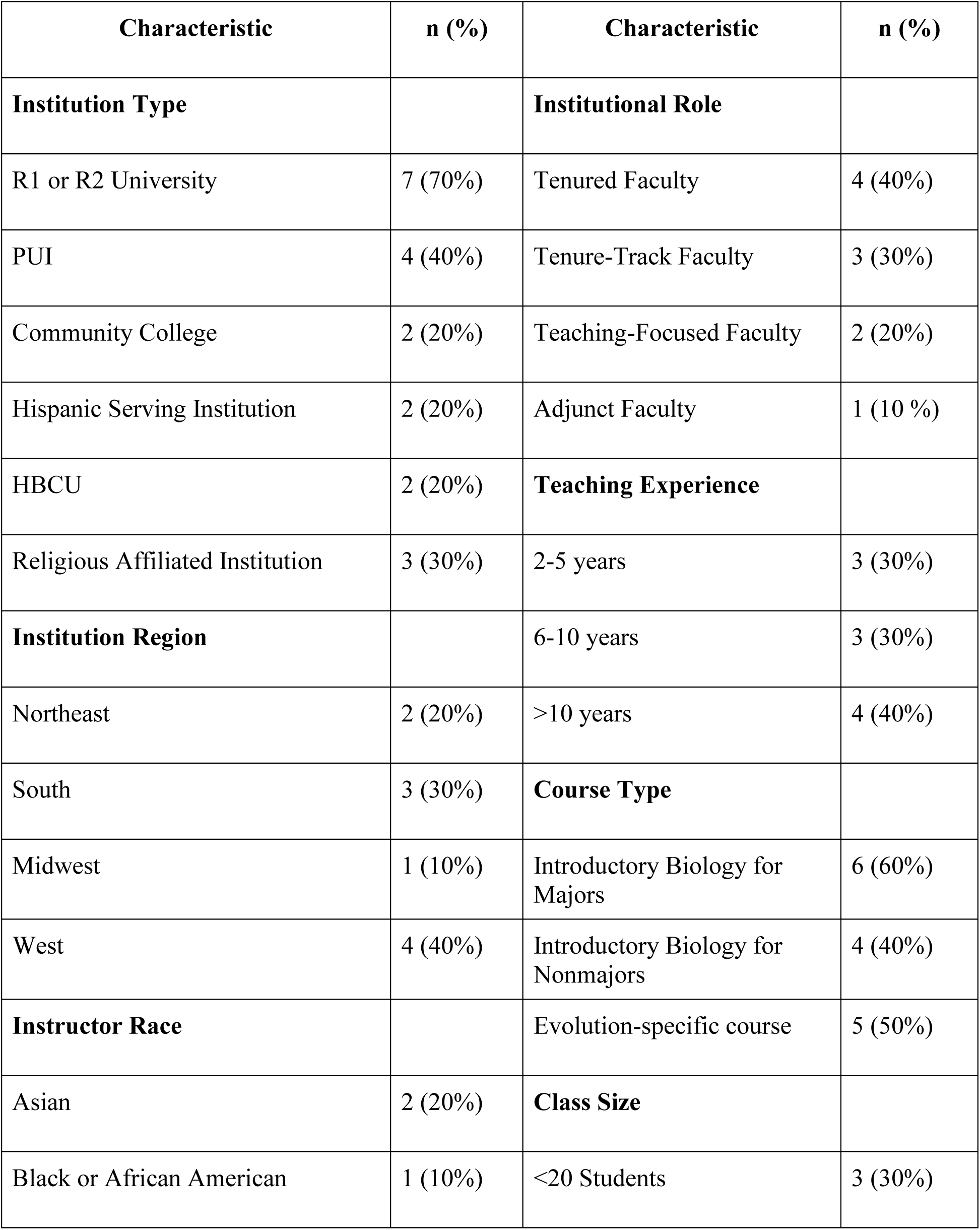
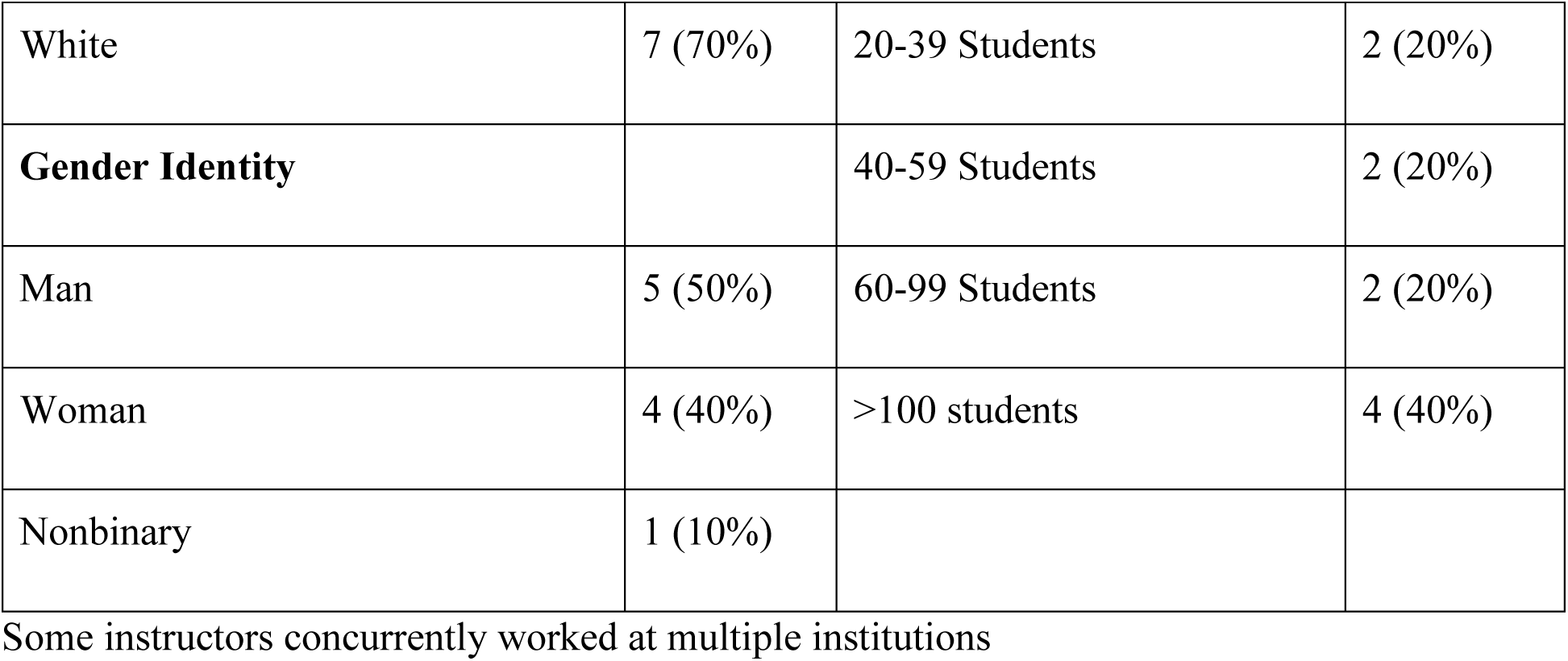
Demographics of Interview Participants (n=10)

### Data Analysis

Interview transcripts were analyzed thematically using an inductive approach (Braun and Clarke, 2006). The first author first read all transcripts in full and generated an initial set of codes based on recurring patterns in instructors’ responses. Subsequently, four additional researchers coded the transcripts, such that each transcript was coded by at least two researchers. Codes were then compared and refined until a minimum of 90% agreement was reached. See Supplemental D for the full codebook.

This research was approved by the St. Mary’s College of Maryland Institutional Review Board #SP26_09. All names represented are pseudonyms.

## RESULTS

### How do instructors feel about game-based learning?

All instructors we interviewed expressed openness to GBL as a tool for evolution instruction, even when they had limited or no experience using video games in their teaching. Several described using related approaches, such as simulations, classroom games, or interactive models. For example, Laura explained that although she did not have instructional experience with video games, she had used HHMI resources and recently designed a board-game-like activity with plastic turtles. She concluded that “anything… that has a game element, and that is interactive in that way would go over really well” with her students. Avery similarly distinguished their use of simulations from video games, but still noted that “there’s a good history of video games being effective teaching tools,” citing Oregon Trail as a classic example of a game used to teach K-12 students about American history. Instructors’ enthusiasm was strongest when they saw games as doing something pedagogically useful beyond adding novelty. Samuel, who stated that he was “not a video game person,” still argued that games could help students because “that’s what they love” and could “help them get motivated, get involved” compared with traditional teaching and PowerPoint lectures. These responses suggest that inexperience with video games among instructors is not likely to preclude their recognition of its potential as a tool for teaching evolution.

### What topics are best for game-based learning?

Instructors’ recommendations for game-based learning were closely tied to their perceptions of student difficulty. The topics they viewed as best suited for gameplay were largely the same topics they described as challenging. Below, we describe each of these five topic areas and examine why instructors viewed them as appropriate for GBL.

#### Population-level reasoning and reasoning across levels

Across interviews, instructors (n=8) described students’ difficulty moving beyond individual organisms to understand evolution as a process that occurs in populations over generations. Daniel identified “thinking about it on a population level rather than the individual level” as one of the major challenges students encounter when learning evolution. James elaborated on this challenge by describing how students must connect multiple levels of biological organization when reasoning about variation and selection. For example, he explained that students were asked to identify sources of phenotypic variation, including “genotypes, environment, or genotype by environment interactions,” and that “it takes a while for them to grasp” that variation comes from these sources. He connected this to broader questions about selection, including “what is the actual unit of selection” and “what’s actually being selected.” Zach similarly described needing to help students understand “the little pieces” of evolution, including how they work and why, before students can move toward broader questions of “what is evolution” and “what does science say it is.”

Instructors also described challenges related to connecting evolutionary processes across scales. Marcus explained that students may understand smaller-scale examples, such as “small tweaks and mutations,” but have more difficulty understanding “how micro evolution build[s] up to become macro evolution.” James also suggested that game-based learning could be useful for representing how selection and drift jointly shape evolutionary change by showing “both selection and drift and their interacting effect on evolution, especially with respect to population size.”

#### Math, probability, and Hardy-Weinberg

Several instructors (n=8) noted that in general, topics related to mathematics and quantitative reasoning were difficult for students to grasp, with Daniel stating that “anything math related is always a bit of a challenge.” Some instructors attributed students’ discomfort with math-related topics to having limited previous math instruction, with Maya stating that many of her students “have had really limited math that has been offered to them” in high school. Marcus shared that many of his students struggle to integrate quantitative concepts into their biology courses, because “biology is underlined by a lot of math,” and students “have a hard time… applying things that they learn in different classes to [biology] class.” In particular, many instructors specified that students struggle to understand Hardy–Weinberg equilibrium. Zach explained that this difficulty may be attributed to the fact that “Hardy–Weinberg is entirely theoretical,” therefore “you can’t really give an example of it.” Daniel shared similar challenges, stating that “a lot of the math and evolution is theoretical,” and “tying in those probabilities and then numbers and math…can be a challenge in a practical sense.”

Some instructors identified concepts related to probability as challenging for students to grasp, with Beth noting that “students struggled with fractions in general,” and Rachel stating that “depending on… who your audience is…probability can be tough.” Other instructors saw probability as something that students can understand given sufficient instruction, with Maya stating that “randomness and probability are concepts that [students] easily can be taught, but have not had the opportunity of being taught yet.” Similarly, when asked how students respond to concepts related to probability, Zach shared that “it just depends if they’ve had stats. More often than not, they have not.” Avery stated that their students seem to generally understand randomness and probability, but they “have a tough time understanding and conceptualizing how those two interact in selection.”

#### Speciation, phylogeny, relatedness

Instructors (n=8) also identified speciation, phylogeny, and relatedness as challenging topics for students. Marcus explained that students often struggle with phylogeny and tree thinking, especially understanding shared ancestry. Laura similarly identified phylogenetics as an area where students have difficulty “identifying… which taxa are more closely related to which other taxa” and “figuring out what constitutes a monophyletic clade.” Instructors also described speciation as challenging because students had difficulty reasoning about how lineages split and change over time. Rachel explained that students struggle with “cladogenesis and anagenesis” and with questions such as “how did something turn into something else?” She further noted that students may understand a lineage splitting when there is a geographic boundary, but have more difficulty when there’s no boundary or when one lineage changes while another remains. Avery also identified speciation as a topic students “struggle with quite a bit,” describing the challenge of condensing the many processes involved in speciation into instruction.

#### Mechanisms of evolution

Instructors (n=8) frequently identified mechanisms of evolution as challenging topics for students, particularly when instruction moved beyond natural selection to include genetic drift, gene flow, mutation, and different types of selection. Daniel described this broader challenge by explaining that many students are familiar with natural selection, but struggle with “all of the other ways in which evolution can occur.” His comment suggests that instructors viewed mechanism-related difficulty as extending beyond whether students could define natural selection; students also needed to recognize that evolutionary change can occur through multiple processes. This pattern was visible in instructors’ descriptions of genetic drift. Beth stated that “students really struggle with genetic drift,” describing it as “the one that’s most confusing to them” among the major evolutionary processes. Similarly, Maya noted that students “struggle with remembering which one’s gene flow [versus] genetic drift.” Instructors also connected mechanism-related challenges to the potential value of visual and interactive representations. Rachel, for example, stated that “genetic drift could be visualized better than the way that most textbooks are doing it,” suggesting that static textbook representations may not fully support the reasoning instructors want students to develop. James made a similar connection to GBL when he suggested that a game could exhibit “both selection and drift and their interacting effect on evolution, especially with respect to population size.”

#### Long time scales and earth history

Instructors (n=6) frequently identified long time scales and Earth history as challenging topics for students. These challenges included reasoning about the age of the Earth, the history of life, and the length of time required for evolutionary change. Daniel described time as difficult to visualize, explaining that students must reason about “how long of a time period” is involved in evolutionary processes and what “10 million years looks like.” He recalled a student asking about the evolution of eukaryotes and noted that although instructors may describe these transitions quickly, they represent “hundreds of millions of years” of change. Rachel similarly emphasized the difficulty of interpreting scale, explaining that students may memorize dates such as when vertebrates evolved, but that “500 million years ago doesn’t really have any meaning” without a way to conceptualize that amount of time. Other instructors described similar challenges when discussing geological time, generational time, and the history of life. Avery emphasized that long time scales are difficult to conceptualize even beyond the classroom, stating that “it’s not…a student problem, it’s an everyone problem.” He described evolutionary and geological time scales as “just unfathomable,” adding, “I don’t actually understand what a million years is like. I don’t understand what 100,000 years is like.” Marcus also described students’ difficulty placing evolutionary events in context, noting that students may try to memorize events such as mass extinctions or rapid diversification without having “a lot of good context for that long history.” Zach described using generational time as one way to discuss evolutionary change, asking students to compare an organism such as a house fly, with a short generation time, to an elephant, with a much longer reproductive cycle. Samuel similarly described Earth history as abstract, explaining that students are asked to “put your mindset in the past, so many years back.”

### Why are these topics difficult for students?

Although instructors were not directly asked to explain why students struggled with specific evolution topics, several offered explanations while discussing challenging content areas. These explanations were unevenly reported: some instructors listed difficult topics without providing a reason, while others described general sources of difficulty that applied across multiple topics. Therefore, we present these categories as emergent themes rather than a comprehensive list of explanations.

A major pattern was the persistence of prior misconceptions about evolutionary processes. Instructors described misconceptions related to the level at which evolution occurs, the role of intention or need, and the relationship between adaptation, selection, and fitness. For example, Rachel emphasized that “the number one thing” students struggle with is that “evolution is not goal directed” and that organisms “didn’t change because they had to.” She noted that even her majors continued to write teleological explanations on exams after repeated instruction. Similarly, Beth described her students interpreting nearly any directional change as adaptation, explaining that once students hear organisms are “evolving in some particular direction,” they “really struggle with thinking about how that can’t be anything except they’re adapting.” Daniel also described repeated difficulty with non-natural selection-based evolution and with helping students to understand that evolution does not always operate through selection alone.

Instructors also described some evolution topics as conceptually difficult even when students were interested and engaged. Avery, for example, explained that selection remained difficult because students often think about it linearly, even though “you can’t tell what selection is going to do” and can often only explain outcomes after the fact. James described a student’s confusion about whether a natural disaster should be understood as drift or selection, noting that students can struggle with the idea that “both are affecting any gene at any time” and that drift and selection may interact depending on population size. Instructors also identified speciation, phylogeny, and tree thinking as difficult because they require students to organize abstract relationships over time.

Several instructors described the language of evolution itself as a barrier. This included both technical vocabulary terms that carry different meanings in everyday, religious, or cultural contexts. Rachel explained that students often enter class with language shaped by high school instruction and non-scientific sources, including social media and news media, which can make it difficult for them to discuss evolution in scientifically accurate ways. She identified fitness as an example, noting that students’ everyday understanding of the word often differs from its meaning in evolutionary biology. Thomas described the word “evolution” as an initial hurdle, explaining that “if you skirt the word and teach the concepts,” he sees “less pushback.” He also noted that students may respond negatively when instructors introduce evolution in a way that feels antagonistic, making the language and framing of instruction important.

Instructors also connected student difficulty to differences in their prior preparation. This included limited exposure to evolution before college, gaps in genetics or mathematics, and varied high school experiences. Marcus described a “massive gradient of preparedness,” with some students coming from strong STEM academies and others taking STEM classes from teachers without STEM backgrounds. He explained that this uneven preparation made it difficult to generalize across students because instructors were trying to “catch them all up and get them on the same page.” Maya similarly connected student difficulty with randomness and probability to limited math preparation, noting that many of her first-year students had “really limited math that has been offered to them.” Beth also described students struggling with Hardy-Weinberg partly because they had difficulty distinguishing “between genotypes and alleles.”

A less frequent but important theme involved religious skepticism and other outside influences that shaped how students approached evolution instruction. Instructors working in religious or regionally conservative contexts described students coming to class with preconceptions that framed evolution as incompatible with faith. Marcus explained that many students in his region had “never heard or thought about this content ever in their life,” making college evolution instruction feel like a “whiplash.” He also described how some students placed “equal weight” on what they learned from family members and what they learned from college instructors. Thomas also described evolution as an “awkward space” for students navigating “evolution and religion or faith,” particularly in the South. Meanwhile, James described using explicit discussion of faith and evolution to help students recognize that some assumptions about evolution were “more cultural than religious.” Importantly, these instructors did not frame religious identity as inherently incompatible with learning evolution; rather, they emphasized that students may need instructional environments that acknowledge students’ preconceived ideas, reduce the perceived conflict, and attempt to create opportunities for students to make sense of the content.

### What would success look like?

When asked what success would look like after implementing an evolution-based video game into their classroom, instructors most frequently described success in terms of improved student learning. Six of the ten instructors pointed to improved assessment performance, including higher exam scores, better response to evolution questions, or gains on pre/post measures. For example, Rachel explained that “getting a better grade on the [test questions] would be really successful,” while Daniel described improvement on persistent areas of difficulty as “a big indicator” that the intervention was working. However, instructors did not define success only as higher scores. Five instructors also emphasized deeper conceptual understanding, as well as an increased ability for students to explain evolutionary mechanisms, accurately use terminology, and reason through new scenarios. Avery, for instance, wanted students to “more accurately describe in their own terminology how a thing works,” while Laura described success as students being able to interpret a new situation or primary literature example and “reason through it effectively.” A smaller number of instructors also described success affectively, including increased excitement, motivation, or openness to learning.

### What concerns do instructors have about implementing a game?

When asked about concerns or hesitancies, instructors were generally open to implementing an evolution video game but emphasized several conditions that would shape integration to their course curriculum. Most concerns centered on whether the game would be instructionally useful, scientifically accurate, accessible, technically feasible, and easy to fit into existing course structures. Instructors wanted the game to be engaging without sacrificing educational value. As Daniel explained, the challenge would be finding “that balance between it being educational enough” while still being “fun and engaging.” Others emphasized the importance of scientific accuracy, noting that gamification could unintentionally introduce misconceptions if evolutionary processes were oversimplified or distorted. Avery, for example, wanted to ensure that the game did not “sacrifice…the true to make something work,” while Laura noted that misconceptions might “sneak in” through the process of gamification. Instructors also raised practical concerns related to accessibility, device compatibility, and time. Rachel emphasized that instructions may be unable to use resources if it is not “fully accessible,” while Beth noted that many students rely on Chromebooks or tablets that may not support more complex games.

## DISCUSSION

In this study, we explored how undergraduate biology instructors evaluated video games as potential tools for evolution education. Instructors were broadly open to game-based learning (GBL), even when they had limited personal or professional experience using video games in their teaching. Their openness was not based only on the assumption that students would find games engaging. Rather, instructors saw games as potentially useful when they could address persistent instructional challenges in evolution, including student misconceptions, difficulty reasoning across populations and time scales, challenges with quantitative reasoning, and tensions related to religion or prior beliefs. The topics instructors viewed as best suited for gameplay were largely the same topics they described as most difficult for students, suggesting that instructors saw GBL as valuable when it could make hard-to-observe evolutionary processes visible, interactive, and connected to learning goals. However, instructors also raised concerns about scientific accuracy, accessibility, feasibility, and curricular alignment. In the following sections, we discuss how these findings extend prior work on evolution education, GBL, and instructor adoption of educational technologies.

### Evolution Learning Challenges as Design Priorities for GBL

Our findings align with prior evolution education literature showing that students struggle with several core concepts instructors identified as promising for GBL Prior work has documented persistent challenges with evolutionary mechanisms and relatedness (Gregory, 2009; Pobiner, 2016). Gregory (2009), for example, argues that natural selection is often poorly understood because students must consider variation within populations, heritability, differential survival and reproduction, and change across generations. Similarly, Pobiner (2016) identifies misconceptions, terminology, cognitive obstacles, and religious worldviews as recurring barriers to accepting and understanding evolution. In our study, instructors identified many of these same areas as both difficult for students and appropriate for GBL. This overlap suggests that instructors’ recommendations were grounded in well-documented challenges in evolution education rather than in a general belief that games are inherently engaging.

Our findings extend this literature by showing how instructors translate known student difficulties into design priorities for instructional games. Prior research has often focused on what students misunderstand, such as the belief that individuals evolve, that evolution is goal-directed, or that evolutionary change is always adaptive or improvement-oriented (Gregory, 2009; Pobiner, 2016; Rice et al., 2015). Instructors in this study identified these same misconceptions, but framed them in terms of what a game could help students see or do. For example, instructors described games as potentially useful for visualizing populations changing over time, representing genetic drift more effectively than textbooks, comparing the effects of drift and selection, modeling lineage splitting, and helping students move across evolutionary or geological time scales. Thus, this study contributes an instructor-centered perspective to the literature by identifying where instructors believe GBL may offer representational advantages over static images, lecture explanations, or isolated classroom examples.

The relationship among identity, religion, and evolution acceptance is complex, with prior work showing that perceived conflict between evolution and religion can strongly shape students’ acceptance of evolution (Barnes et al., 2021). Although GBL shows promise for helping students visualize and reason through evolutionary processes, students may benefit from instructional supports that address broad cultural and identity-based aspects of learning evolution. For example, conflict-reducing practices such as acknowledging students concerns, teaching the bounded and agnostic nature of science, and affirming students’ autonomy may help reduce perceived conflict and create more supportive learning environments (Barnes and Brownell, 2026, 2017; Bertka et al., 2019). Recent work by Rowland-Schaefer and colleagues (2026) further stresses the need for these supports, showing that college biology instructors vary in whether they treat evolution acceptance as an instructional goal and that many do not discuss the relationship between evolution and religion when teaching evolution. In the context of GBL, this suggests that evolution-focused games may be most effective when paired with instructor-led discussion, reflection, or debriefing that helps students connect gameplay to scientific explanations while also acknowledging the prior beliefs and concerns they may bring to the classroom.

### Instructor Perspectives on Game-Based Learning

While there is broad agreement that GBL can be an effective pedagogical tool, the effectiveness of a game depends on design quality and pedagogical alignment (Clark et al., 2016; Linderoth, 2012; Subhash and Cudney, 2018). Developing a successful educational game therefore requires multiple forms of expertise, including content knowledge, pedagogical knowledge, and game design experience (Linderoth and Sjöblom, 2019). However, these forms of expertise are not always held by the same people. Instructors may have the disciplinary and pedagogical knowledge needed to ensure curricular alignment, but they rarely have the time, training, or institutional support required to build games themselves (Dimitriadou et al., 2021). Conversely, game developers may be highly skilled in designing engaging gameplay, but may prioritize creative or design elements unless educational goals are clearly integrated into the development process (Linderoth and Sjöblom, 2019). Prior research on multidisciplinary educational game development teams has shown that collaboration between content experts and game developers can be challenging because of differences in vocabulary, expectations, timelines, and information-sharing practices (Korhonen et al., 2017). However, research on GBL also supports the involvement of instructors in the development of games they will use in their classrooms (Vlachopoulos and Makri, 2017).

Our findings support the importance of instructor involvement in educational game design. Although prior studies suggest that personal and professional inexperience with video games can be a barrier to instructors viewing GBL as a valid instructional tool (Bourgonjon et al., 2013), instructors in this study were broadly open to the possibility that a video game could support evolution learning. This openness is notable because many instructors did not describe themselves as gamers or as having extensive experience with video games in the classroom. Instead, they evaluated GBL through the lens of instructional usefulness. Instructors were most enthusiastic when they could imagine a game helping students visualize populations changing over time, compare evolutionary mechanisms, or interact with processes that are otherwise difficult to observe and even more difficult to visually and conceptually connect (e.g. molecular processes and population-level selective processes). This suggests that instructor adoption of GBL may depend less on whether instructors personally value video games and more on whether they see a clear connection between the game, the learning challenge, and their course goals.

Instructors’ concerns also provide important guidance for educational game design. Some concerns were practical, including limited class time, technical feasibility, student access, and regional or institutional restrictions on instructional materials. Other concerns were pedagogical, particularly the possibility that “gamification” could sacrifice scientific accuracy or reinforce misconceptions. These concerns are consistent with prior arguments that educational games must be designed carefully to avoid prioritizing engagement at the expense of learning (Linderoth and Sjöblom, 2019). However, the concerns raised by instructors were also highly actionable. For example, accessibility concerns can guide decisions about contrast, captions, navigation, and device requirements, while concerns about misconceptions can guide decisions about how evolutionary mechanisms are represented in the game. Rather than viewing instructor concerns as barriers to adoption, these concerns can be treated as design criteria for creating games that are scientifically accurate, accessible, and usable in real classrooms.

### Implications and Future Directions

The findings from this study have direct implications for future development of evolution-focused video games and other interactive instructional tools. Instructors repeatedly identified the need for tools that represent evolution as a population-level process unfolding over time. Therefore, games designed for evolution education should avoid centering only on individual organisms as the unit of evolutionary change unless gameplay clearly connects individual variation, survival, reproduction, and inheritance to population-level outcomes. Similarly, because instructors identified genetic drift, selection, gene flow, speciation, probability, and long time scales as challenging topics, future games may be useful when they allow students to manipulate conditions and observe how evolutionary processes operate across generations. Interview feedback suggests that students may benefit from connecting elements across the scope of evolution by connecting molecular processes to individuals, individuals to populations, populations to species, and so on.

Future work should examine whether evolution games support learning in the specific areas instructors identified as challenging. For example, studies could assess whether gameplay improves students’ ability to reason across individuals and populations, distinguish among evolutionary mechanisms, interpret lineage splitting, or conceptualize deep time. These outcomes could be examined using a combination of validated concept inventories, pre/post assessments, and open-ended response questions aligned with the specific concepts addressed in the game. It will also be important to examine whether students can transfer understanding beyond the game context. Students may succeed within a game by manipulating traits, populations, or environments, but additional research is needed to determine whether they can explain evolutionary processes accurately using biological language after gameplay.

Future research should also examine how instructors implement evolution games in different classroom contexts. Instructor framing may be important for topics that are both conceptually difficult and culturally sensitive. A game may help students visualize genetic drift or lineage splitting, but it may not address perceived conflict between evolution and religion unless instructors intentionally create space for students to engage with the content in a respectful and scientifically grounded way. Therefore, future studies should examine not only whether an evolution game improves learning, but how it works when embedded within different instructional approaches, student populations, and institutional contexts.

Finally, this study supports the need for collaborative design models that include instructors, evolution education researchers, students, and game developers. Instructors in this study identified both opportunities and risks that may not be immediately visible to game designers working outside of classroom contexts. Their insights can help ensure that evolution games are not only engaging, but also accurate, accessible, and aligned with the learning goals of biology courses. More broadly, our findings suggest that instructor perspectives are essential for designing educational technologies that respond to real instructional challenges rather than assuming that engagement alone will improve learning.

## ACKNOWLEDGEMENTS

We are grateful to the evolution instructors who generously shared their time, teaching experiences, and perspectives on game-based learning. Their insights were essential to this work. We also acknowledge funding support from the National Science Foundation to A.E.B. and J.E.P. (NSF-IUSE-2439101).

## Author Contributions

Josie Otto: Data curation, Formal analysis, Investigation, Methodology, Resources, Writing – original draft, Writing – review & editing, Supervision.

Lauren Goulet: Investigation, Writing – original draft.

Brooke Kopack Ware: Investigation, Writing – original draft.

Hannah Lowry: Investigation, Writing – original draft.

Aeryn-Elayna Miller: Investigation, Writing – original draft.

Jake D. Botello: Conceptualization, Formal analysis, Funding acquisition, Writing – review & editing.

Jenna E. Pruett: Conceptualization, Funding acquisition, Supervision, Writing – original draft.

Abby Beatty: Conceptualization, Data curation, Formal analysis, Funding acquisition, Methodology, Project administration, Supervision, Visualization, Writing – review & editing.

## Generative AI Use

The authors report generative AI was not used in their research or preparation of this manuscript

## Supplemental A. Instructor Screening Survey

Thank you for your interest in participating in our NSF-funded research study examining the use of an educational video game to support undergraduate evolution education.

This brief screening survey (approximately <5 minutes) will help the research team identify eligible participants and select a diverse group of instructors for expert interviews. Completion of this survey does not guarantee selection.

1. Are you currently teaching or have you recently taught undergraduate students in biology or a related STEM field?

☐ Yes

☐ No **<<END SURVEY and display the following message: Based on your response, you are not eligible to continue with the survey at this time. This study is designed for instructors who currently teach or have recently taught undergraduate biology or related STEM courses that include evolutionary concepts. We appreciate your time and willingness to participate.>>**

2. Does your teaching include evolution or evolutionary concepts (e.g., natural selection, gene flow, mutation, drift, common ancestry)?

☐ Yes

☐ No **<<END SURVEY and display the following message: Based on your response, you are not eligible to continue with the survey at this time. This study is designed for instructors who currently teach or have recently taught undergraduate biology or related STEM courses that include evolutionary concepts. We appreciate your time and willingness to participate.>>**

3. Institution Type

(Select all that apply)

☐ Community College

☐ Primarily Undergraduate Institution (PUI)

☐ Master’s-granting institution

☐ Research-intensive university (R1 or R2)

☐ Historically Black College or University (HBCU)

☐ Hispanic-Serving Institution (HSI)

☐ Religious-affiliated institution

☐ Tribal College or University

☐ Minority-Serving Institution (other)

☐ Other (please specify):

4. Geographic Location of Institution

☐ Northeast

☐ Midwest

☐ South

☐ West

☐ Outside the United States

5. What is your current instructional role?

☐ Tenured faculty

☐ Tenure-track faculty

☐ Lecturer / Instructor

☐ Teaching-focused faculty

☐ Adjunct faculty

☐ Graduate student instructor

☐ Other (please specify)

6. How many years have you been teaching at the undergraduate level?

☐ Less than 2 years

☐ 2–5 years

☐ 6–10 years

☐ More than 10 years

7. Which course levels do you currently teach?

(Select all that apply)

☐ Introductory biology (majors)

☐ Introductory biology (non-majors)

☐ Upper-division biology

☐ Evolution-specific course

☐ Other (please specify)

8. Approximate enrollment in the course(s) where you teach evolution:

☐ Fewer than 20 students

☐ 20–39 students

☐ 40–59 students

☐ 60–99 students

☐ 100+ students

9. How do you describe your gender identity?

☐ Woman

☐ Man

☐ Nonbinary / gender diverse

☐ Prefer to self-describe:

☐ Prefer not to say

10. How do you describe your race and/or ethnicity?

☐ American Indian or Alaska Native

☐ Asian

☐ Black or African American

☐ Hispanic or Latino/a/e

☐ Native Hawaiian or Other Pacific Islander

☐ White

☐ Multiracial

☐ Prefer to self-describe:

☐ Prefer not to say

11. How would you describe your background with video games? (Select all that apply.)

☐ I regularly play video games recreationally

☐ I occasionally play video games recreationally

☐ I have played video games in the past but not recently

☐ I do not play video games and have little to no experience with them

☐ I have used video games in an educational or instructional context

☐ I have professional experience related to video games (e.g., development, research, design, industry)

☐ Other (please describe):

12. **If selected, what is the best email address to contact you?**

**<<END SURVEY and display the following message: Your responses indicate that you may be eligible to participate in this study. The research team will review all completed surveys, and if you are selected, you will be contacted with additional information about the next steps.**

**We sincerely appreciate your time and contribution to this research.>>**

## Supplemental B. Instructor Interview Screening Survey Demographics (n=39)

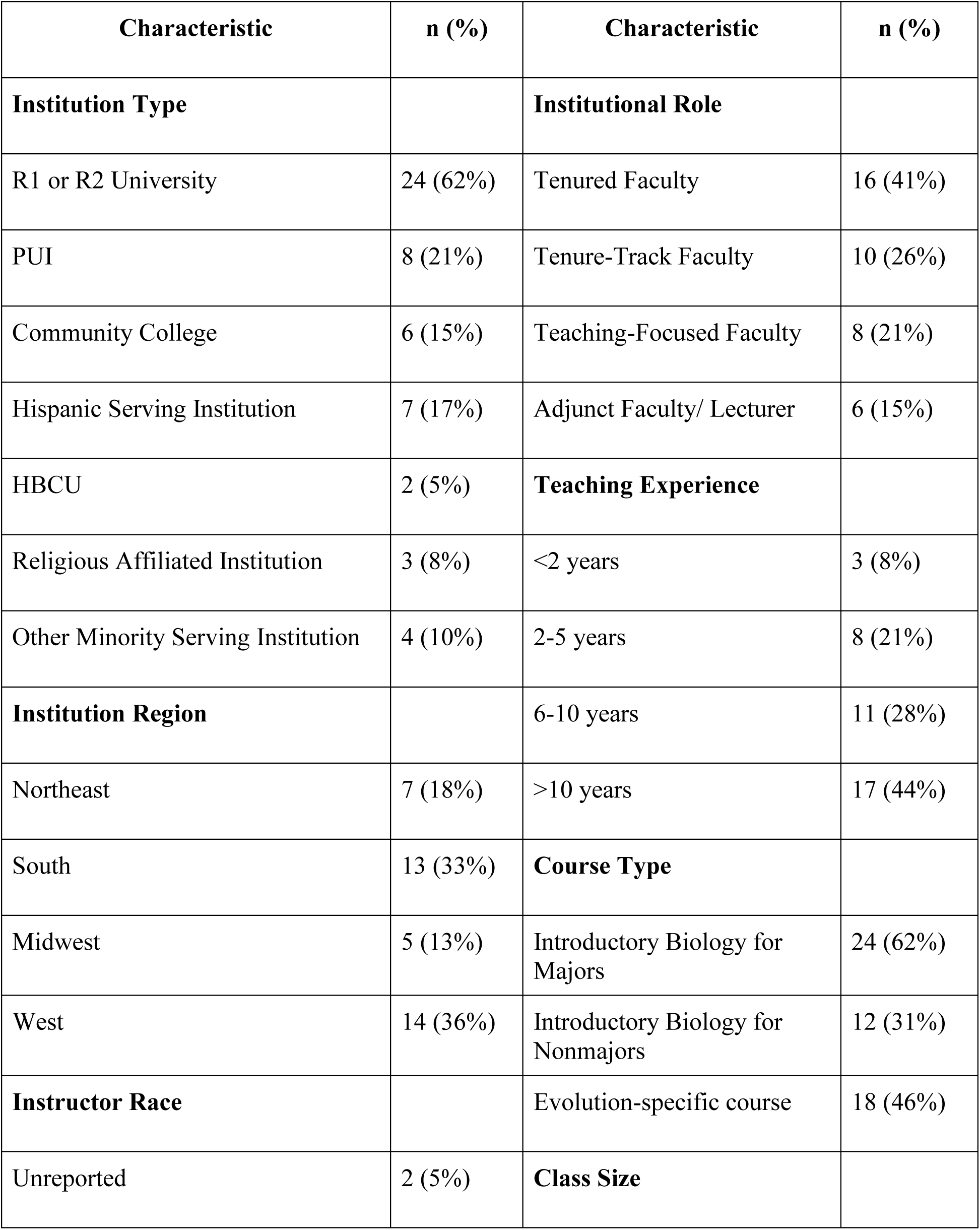
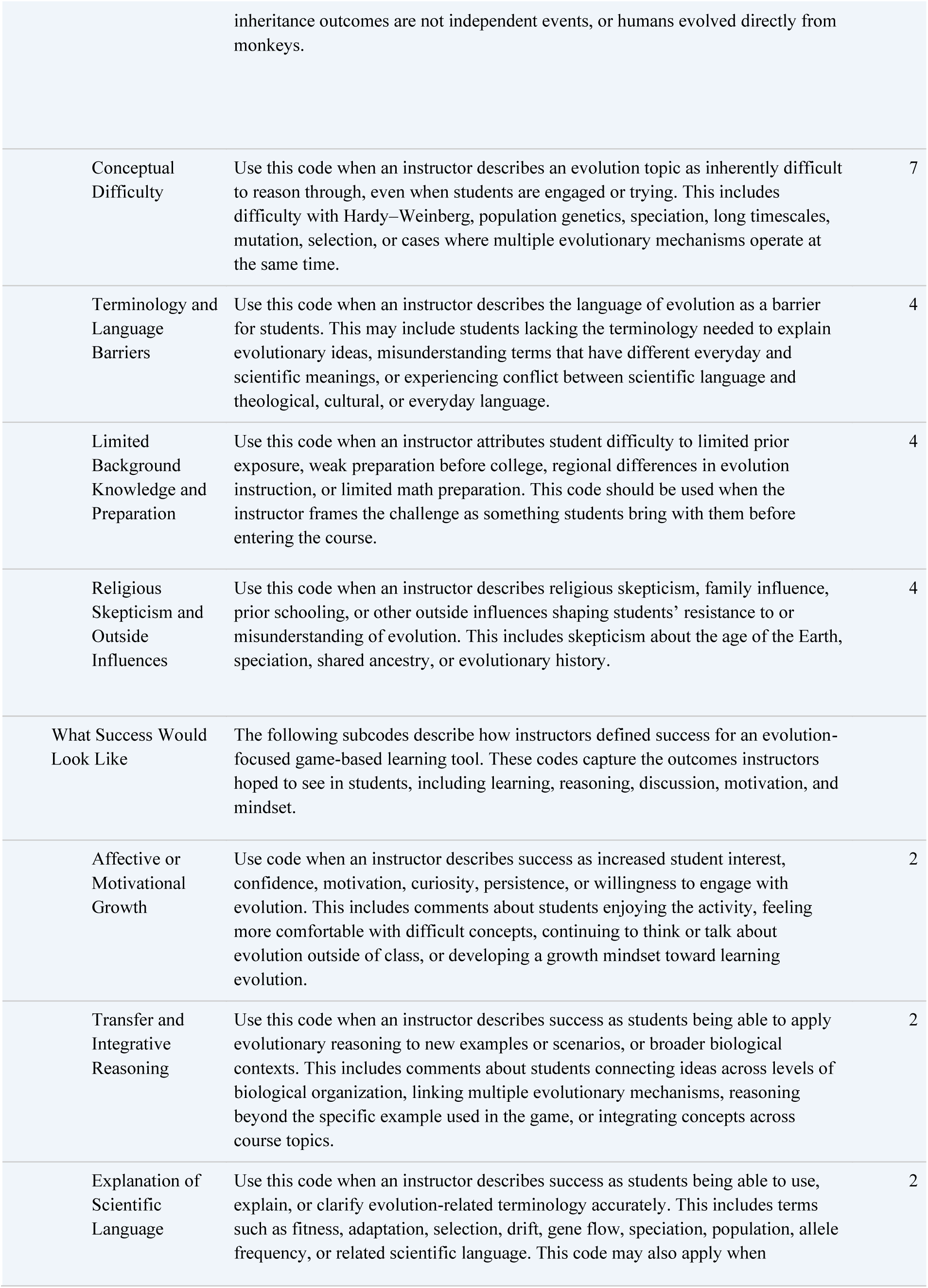
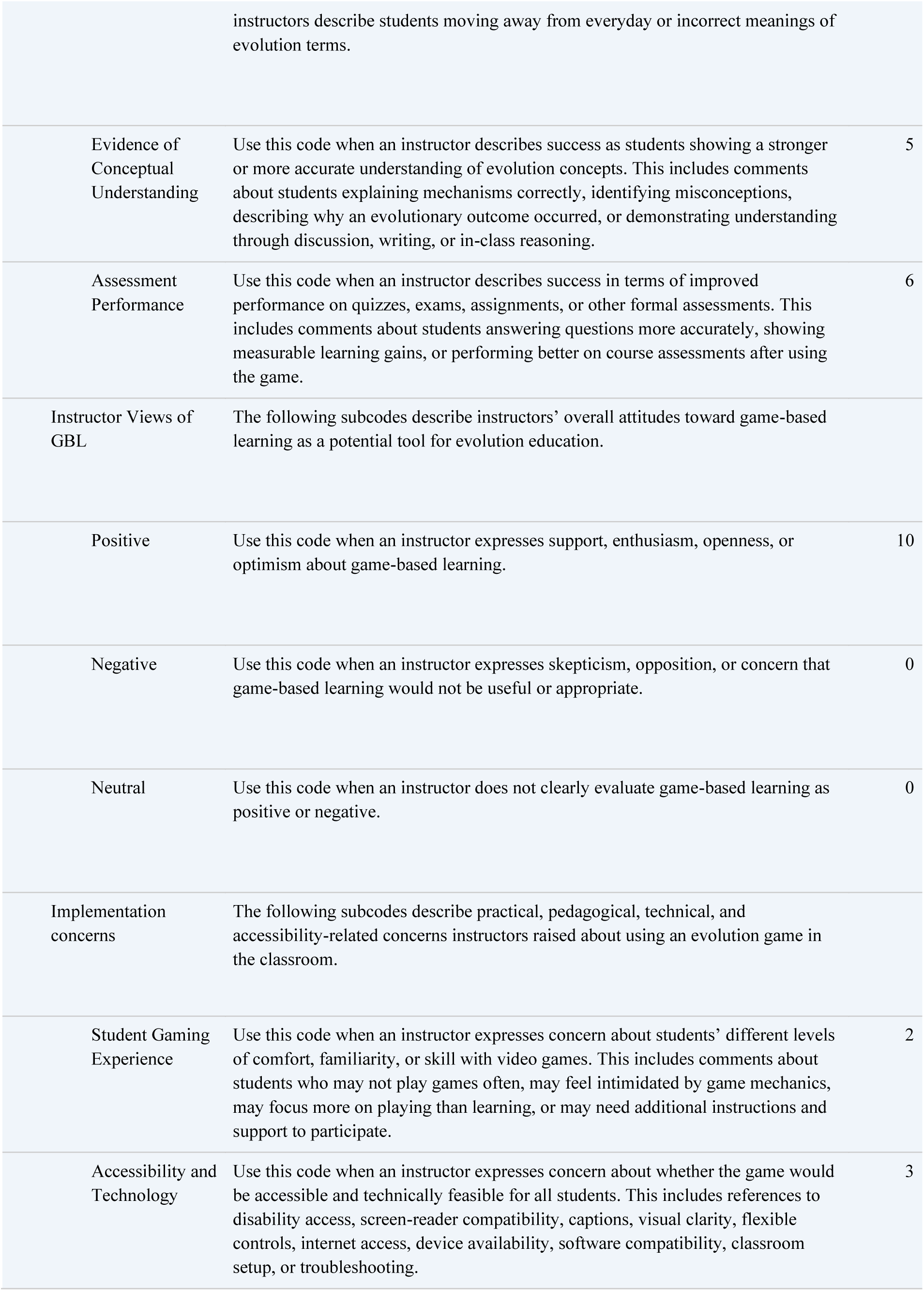
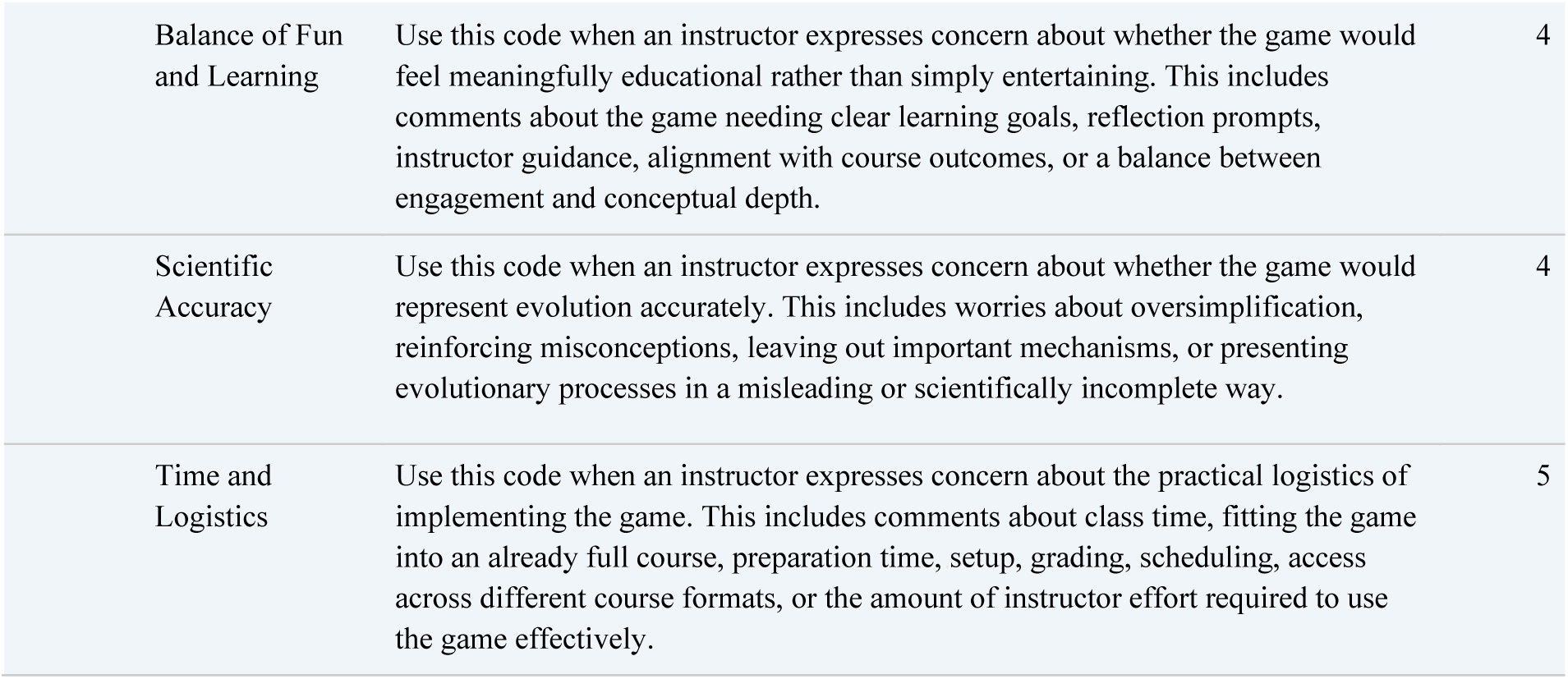

## Supplemental C. Instructor Interview Script and Questions

Hi [Name], thank you so much for taking the time to meet with me today.

My name is [Name], and I’m part of the research team working on an NSF-funded project focused on evolution education and game-based learning. *[If other team members are present*: I am joined today by [Name(s)]. We really appreciate you sharing your time and expertise with us.

Before we begin, I want to briefly explain the purpose of today’s interview.

We’re conducting interviews with biology instructors to better understand how evolution is taught, where students tend to struggle, and how instructional tools, specifically an educational video game, might support student learning and engagement.

Your responses will help inform the design and implementation of this tool. There are no right or wrong answers - we’re interested in your experiences and perspectives as an instructor.

As we go through the interview, we’ll also post each question in the chat so you can reference it as we talk. You’re welcome to look back at the questions at any time, and we can always pause or revisit anything you’d like to expand on.

You received and signed a consent form prior to this interview. I want to briefly review a few key points before we continue.

Your participation in this interview is voluntary, and you may skip any question or stop the interview at any time without penalty.

As indicated in the consent form, this interview will be audio-recorded so we can accurately capture your responses. The recording will be transcribed, and any identifying information will be removed from the transcript. Only the research team will have access to the recordings and transcripts.

Before we begin, do you have any questions about the consent form or the interview process?

For the record, can you please confirm that you consent to participate in this interview and to being audio-recorded?

Thank you. I’ll start the recording now.

1. To start, could you briefly describe the courses you teach that include evolution content and the types of students typically enrolled?
2. Thinking back to your own education, when were you first introduced to evolution (e.g., middle school, high school, undergraduate) and what has your evolution education consisted of throughout college?

a. How would you describe your earliest instructor’s approach or perspective on evolution, and in what ways, if any, has that experience influenced how you teach evolution today?
3. Based on your teaching experience, what aspects of evolution do students find most challenging to understand (e.g., population genetics, fitness, mutation, gene flow, natural selection, genetic drift, nonrandom mating, hardy weinberg, selection, speciation, etc.)?

a. Can you describe specific concepts or ideas that students struggle with repeatedly?
b. Are there challenges that seem to persist even after multiple forms of instruction?
4. Which aspects of evolution or genetics do students seem to have the hardest time visualizing or reasoning through?

a. Do students tend to struggle more with thinking at the genetic, individual level or the population level? -
b. How do students respond to concepts involving randomness, probability, or long time scales?
5. How do students typically respond emotionally or motivationally when learning about evolution? -

a. What signs do you notice when students are engaged versus disengaged? *And is this evident with any particular topics or language/phrases?*
b. Are there moments when students appear uncomfortable or resistant, and how does that show up? *And is this evident with any particular topics or language/phrases?*
6. Do you ever discuss concepts related to god, the origins of life, or metaphysics in the courses where you teach evolution? If so, tell us how you weave them into your curriculum. - ANSWERED
7. How do students’ prior beliefs or backgrounds appear to shape their engagement with evolution content?

a. How do students talk about the relationship between evolution and their personal beliefs, if at all?
b. Are there differences you notice across course levels or student populations?
8. What approaches have you found most effective for helping students understand difficult concepts in evolution?

a. Can you describe an approach that has worked particularly well and why you think it was effective?
b. Are there strategies you rely on when students are not responding to more traditional instruction?
9. How do you balance maintaining scientific accuracy with supporting inclusive classroom discussions?
10. When teaching evolution, how do you decide which concepts to emphasize most or to reinforce?
11. Can you explain your current experience with video games recreationally and professionally?
12. In what ways do you think a video game could support students’ learning or engagement with evolution?

a. What aspects of evolution might benefit most from a game-based approach?
b. Is there anything in particular that you’d want students to see or experience?
c. Are there limitations or concerns you would have about using a video game in this context?
13. How much control should students have over decisions compared to being guided through processes?
14. After students engage with a learning tool like this, what changes would signal success to you?

a. What changes would you hope to see in students’ understanding of evolution?
15. Is there anything about your teaching context or student population that you think would be important for us to consider when designing or implementing this game?
16. When the game is finished, would you be interested in testing the game in your classroom?

1. That’s all the questions I have. Thank you again for sharing your insights - they’re incredibly valuable to this project.

If you have any questions after today or think of anything else you’d like to add, please feel free to reach out.

We’ll follow up with next steps and compensation information shortly. Thanks again for your time and expertise.

## Supplemental D. Instructor Interview Codebook

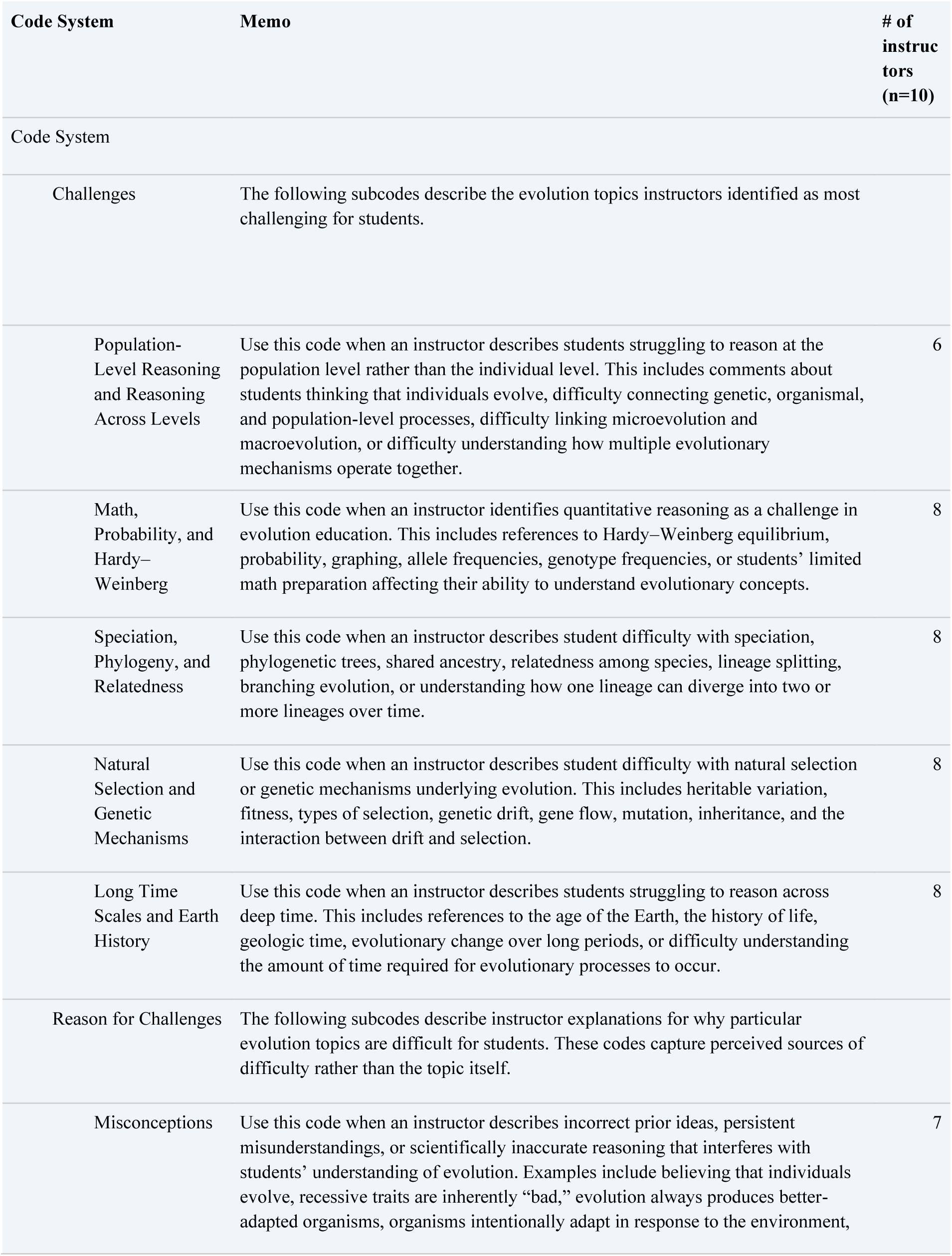

## References

Adams, S., Ajlouni, J., Dennis, A., Rossi, M., 2011. Adventure club games presskit—evolution games [WWW Document]. URL http://adventureclubgames.com/press/sheet.php?p=Evolution_Games

Al-Elq, A., 2010. Simulation-based medical teaching and learning. J. Fam. Community Med. 17, 35. 10.4103/1319-1683.68787

American Association for the Advancement of Science, 2011. Vision and change in undergraduate biology education: a view for the 21st century.

Barnes, M.E., Aini, R.Q., Collins, J.P., Dunk, R.D.P., Holt, E.A., Jensen, J., Klein, J.R., Misheva, T., Nadelson, L.S., Reiss, M.J., Romine, W.L., Shtulman, A., Townley, A.L., Wiles, J.R., Zheng, Y., Brownell, S.E., 2024. Evaluating the current state of evolution acceptance instruments: a research coordination network meeting report. Evol. Educ. Outreach 17, 1. 10.1186/s12052-024-00194-0

Barnes, M.E., Brownell, S.E., 2026. Teaching undergraduates evolution: 10 years of research on conflict-reducing practices—and resistance to them. BioScience biag039. 10.1093/biosci/biag039

Barnes, M.E., Brownell, S.E., 2017. A Call to Use Cultural Competence When Teaching Evolution to Religious College Students: Introducing Religious Cultural Competence in Evolution Education (ReCCEE). CBE—Life Sci. Educ. 16, es4. 10.1187/cbe.17-04-0062

Barnes, M.E., Riley, R., Bowen, C., Cala, J., Brownell, S.E., 2022. Community College Student Understanding and Perceptions of Evolution. CBE—Life Sci. Educ. 21, ar46. 10.1187/cbe.21-09-0229

Barnes, M.E., Supriya, K., Zheng, Y., Roberts, J.A., Brownell, S.E., 2021. A New Measure of Students’ Perceived Conflict between Evolution and Religion (PCoRE) Is a Stronger Predictor of Evolution Acceptance than Understanding or Religiosity. CBE—Life Sci. Educ. 20, ar42. 10.1187/cbe.21-02-0024

Bean, T.E., Sinatra, G.M., Schrader, P.G., 2010. Spore: Spawning Evolutionary Misconceptions? J. Sci. Educ. Technol. 19, 409–414.

Bertka, C.M., Pobiner, B., Beardsley, P., Watson, W.A., 2019. Acknowledging students’ concerns about evolution: a proactive teaching strategy. Evol. Educ. Outreach 12, 3. 10.1186/s12052-019-0095-0

Bourgonjon, J., De Grove, F., De Smet, C., Van Looy, J., Soetaert, R., Valcke, M., 2013. Acceptance of game-based learning by secondary school teachers. Comput. Educ. 67, 21–35.

Braun, V., Clarke, V., 2006. Using thematic analysis in psychology. Qual. Res. Psychol. 3, 77–101. 10.1191/1478088706qp063oa

Brownell, S.E., Freeman, S., Wenderoth, M.P., Crowe, A.J., 2014. BioCore Guide: A Tool for Interpreting the Core Concepts of Vision and Change for Biology Majors. CBE—Life Sci. Educ. 13, 200–211. 10.1187/cbe.13-12-0233

Clark, D.B., Tanner-Smith, E.E., Killingsworth, S.S., 2016. Digital Games, Design, and Learning: A Systematic Review and Meta-Analysis. Rev. Educ. Res. 86, 79–122. 10.3102/0034654315582065

Dankbaar, M.E., Alsma, J., Jansen, E.E., van Merrienboer, J.J., van Saase, J.L., Schuit, S.C., 2016. An experimental study on the effects of a simulation game on students’ clinical cognitive skills and motivation. Adv. Health Sci. Educ. 21, 505–521.

Dimitriadou, A., Djafarova, N., Turetken, O., Verkuyl, M., Ferworn, A., 2021. Challenges in serious game design and development: Educators’ experiences. Simul. Gaming 52, 132–152.

Dobzhansky, T., 1964. Biology, molecular and organismic. Am. Zool. 4, 443–452.

Donald, M.M., 2025. The Impact of PhET Simulations on the Teaching and Learning of Natural Sciences in Grade 7 Classrooms (Master’s Thesis). University of South Africa (South Africa).

Furrow, R.E., Hsu, J.L., 2019. Concept inventories as a resource for teaching evolution. Evol. Educ. Outreach 12, 2. 10.1186/s12052-018-0092-8

Gilbert, J.K., Osborne, R., 1980. The use of models in science and science teaching. Eur. J. Sci. Educ. 2, 3–13.

Gini, F., Bassanelli, S., Bonetti, F., Mogavi, R.H., Bucchiarione, A., Marconi, A., 2025. The role and scope of gamification in education: A scientometric literature review. Acta Psychol. (Amst.) 259, 105418.

Gregory, T.R., 2009. Understanding Natural Selection: Essential Concepts and Common Misconceptions. Evol. Educ. Outreach 2, 156–175. 10.1007/s12052-009-0128-1

Harahap, F.S., Susetyarini, E., Purwanti, E., Fitri, S., Rukman, N.K., Pohan, H.M., 2025. PhET simulation in education: A bibliometric analysis of the Scopus database. Res. Dev. Educ. RaDEn 5, 555–570. 10.22219/raden.v5i1.40504

Heeter, C., Starr, B., 2012. DNA roulette: understanding genetics and genetic testing through gaming [WWW Document].

Korhonen, T., Halonen, R., Ravelin, T., Kemppainen, J., Koskela, K., 2017. A multidisciplinary approach to serious game development in the health sector, in: Proceedings of the 11th Mediterranean Conference on Information Systems MCIS 2017. September 4-5, 2017, Genoa, Italy. Association for Information Systems.

Leith, A.P., Ratan, R.A., Wohn, D.Y., 2016. The (De-)evolution of Evolution Games: A Content Analysis of the Representation of Evolution Through Natural Selection in Digital Games. J. Sci. Educ. Technol. 25, 655–664. 10.1007/s10956-016-9620-x

Linderoth, J., 2012. Why gamers don’t learn more: An ecological approach to games as learning environments. J. Gaming Virtual Worlds 4, 45–62.

Linderoth, J., Sjöblom, B., 2019. Being an educator and game developer: The role of pedagogical content knowledge in non-commercial serious games production. Simul. Gaming 50, 771–788.

Misheva, T., Coburn, K.N., Wiles, J.R., 2026. Are They Cousins? Exploring University Students’ Acceptance of Common Ancestry in Evolution. CBE—Life Sci. Educ. 25, ar1. 10.1187/cbe.25-07-0155

Nkhoma, M., Calbeto, J., Sriratanaviriyakul, N., Muang, T., Ha Tran, Q., Kim Cao, T., 2014. Towards an understanding of real-time continuous feedback from simulation games. Interact. Technol. Smart Educ. 11, 45–62. 10.1108/ITSE-03-2013-0005

Owolabi, J.O., Gardner, K., Agboola, R., Yesudas, R.R., Shaw, J.H., 2025. Use of simulation for teaching biomedical sciences to undergraduate medical students- a scoping review. BMC Med. Educ. 25, 1259. 10.1186/s12909-025-07819-y

Plass, J.L., Homer, B.D., Kinzer, C.K., 2015. Foundations of Game-Based Learning. Educ. Psychol. 50, 258–283. 10.1080/00461520.2015.1122533

Pobiner, B., 2016. Accepting, understanding, teaching, and learning (human) evolution: Obstacles and opportunities. Am. J. Phys. Anthropol. 159, 232–274. 10.1002/ajpa.22910

Poli, D., Berenotto, C., Blankenship, S., Piatkowski, B., Bader, G.A., Poore, M., 2012. Bringing Evolution to a Technological Generation: A Case Study with the Video Game SPORE. Am. Biol. Teach. 74, 100–103. 10.1525/abt.2012.74.2.7

Ratinho, E., Martins, C., 2023. The role of gamified learning strategies in student’s motivation in high school and higher education: A systematic review. Heliyon 9. 10.1016/j.heliyon.2023.e19033

Rice, J.W., Clough, M.P., Olson, J.K., Adams, D.C., Colbert, J.T., 2015. University faculty and their knowledge & acceptance of biological evolution. Evol. Educ. Outreach 8, 8. 10.1186/s12052-015-0036-5

Rowland-Schaefer, E.G., Edwards, B.A., Crowe, A.J., Barnes, M.E., Brownell, S.E., 2026. Challenges to evolution as a core concept in college biology: Silence on religion and conflicting goals for acceptance. BioScience biag072. 10.1093/biosci/biag072

Rutten, N., Van Joolingen, W.R., Van Der Veen, J.T., 2012. The learning effects of computer simulations in science education. Comput. Educ. 58, 136–153.

Shaffer, D.W., Squire, K.R., Halverson, R., Gee, J.P., 2005. Video games and the future of learning. Phi Delta Kappan 87, 105–111.

Snow, E.L., Allen, L.K., Jacovina, M.E., McNamara, D.S., 2015. Does agency matter?: Exploring the impact of controlled behaviors within a game-based environment. Comput. Educ. 82, 378–392. 10.1016/j.compedu.2014.12.011

Su, M., Cho, J.Y., Chi, M.T., Boucher, N., Vanbibber, B., 2021. Designing simulation module to diagnose misconceptions in learning natural selection, in: Proceedings of the 15th International Conference of the Learning Sciences-ICLS 2021. International Society of the Learning Sciences.

Subhash, S., Cudney, E.A., 2018. Gamified learning in higher education: A systematic review of the literature. Comput. Hum. Behav. 87, 192–206.

Vlachopoulos, D., Makri, A., 2017. The effect of games and simulations on higher education: a systematic literature review. Int. J. Educ. Technol. High. Educ. 14, 22. 10.1186/s41239-017-0062-1

